# Profiling and Functional Analysis of Urinary Exosomal MicroRNAs in Pregnant Women with Systemic Lupus Erythematosus

**DOI:** 10.1101/2025.07.29.667355

**Authors:** Dong Li, Anna Li, Linghong Liu, Yuan Liu

## Abstract

**Background:** Pregnancy in Systemic Lupus Erythematosus (pSLE) is high-risk, necessitating non-invasive biomarkers for monitoring and predicting complications. Urinary exosomes, containing miRNAs, offer a promising source reflecting systemic and renal states, yet their profile in late gestation pSLE is less studied.

**Objective:** This study aimed to investigate the profile of urinary exosomal miRNAs in pregnant women with SLE during late gestation compared to healthy pregnant controls and to explore their potential biological roles and pathways.

**Methods:** Urinary exosomes were isolated from 6 pSLE patients and 5 controls. Exosomes were characterized, and miRNAs were sequenced. Bioinformatics analyses, including target gene prediction and functional enrichment (GO, KEGG), were performed.

**Results:** Exosomes were successfully isolated. Sequencing identified 29 significantly downregulated miRNAs in pSLE exosomes. Enrichment analysis of these miRNAs and their target genes indicated involvement in metabolic and immune pathways, virus infection, and cellular senescence. Target genes also enriched in protein stability and transport, and high-level analysis showed broader roles in metabolic, immune, and cellular maintenance/stress processes. Targets of top differentially expressed miRNAs linked to mRNA regulation, AMPK, and senescence pathways.

**Conclusion:** Our findings demonstrate that urinary exosomal miRNAs are significantly altered in pregnant women with SLE during late gestation. The functional enrichment analysis suggests their target genes are involved in critical biological processes relevant to pSLE pathophysiology and pregnancy complications, including metabolism, immunity, viral response, senescence, and cellular maintenance. This highlights the potential of urinary exosomal miRNAs as non-invasive biomarkers for monitoring and predicting risks in this high-risk population.

## 1. Introduction

Systemic Lupus Erythematosus (SLE) is a chronic, complex autoimmune disease characterized by systemic inflammation that can affect virtually any organ system, leading to a wide spectrum of clinical manifestations [1]. The disease primarily affects women of childbearing age, making pregnancy a significant consideration for many patients. Pregnancy in women with SLE (pSLE) is considered high-risk, associated with increased rates of maternal complications, such as disease flares, lupus nephritis, preeclampsia, and thromboembolic events, as well as adverse fetal outcomes, including spontaneous abortion, intrauterine growth restriction, prematurity, and neonatal lupus [2, 3]. Managing pSLE is challenging, requiring careful monitoring of disease activity and potential complications throughout gestation to ensure optimal outcomes for both mother and fetus. Accurate, timely, and non-invasive methods for assessing disease status and predicting risks during pregnancy are highly needed.

Extracellular vesicles (EVs), including exosomes, microvesicles, and apoptotic bodies, are nano-sized lipid bilayer-bound vesicles released by various cell types [4]. EVs are present in almost all biological fluids, including blood, urine, amniotic fluid, and breast milk, and play crucial roles in intercellular communication by transferring their diverse cargo, which includes proteins, lipids, and nucleic acids, particularly microRNAs (miRNAs) [5]. miRNAs are small, non-coding RNA molecules that regulate gene expression at the post-transcriptional level and are involved in a multitude of biological processes, including immune responses, inflammation, and organ development [6].

Urinary exosomes are derived from cells throughout the nephron and urinary tract, offering a valuable and non-invasive window into the physiological and pathological state of the kidney and potentially reflecting systemic conditions affecting these organs [7]. Given the high prevalence of lupus nephritis in SLE, and its potential to flare or complicate pregnancy in pSLE patients, urinary exosomes and their molecular contents, such as miRNAs, hold significant promise as potential biomarkers for monitoring disease activity or predicting complications, particularly in the context of pSLE.

While studies have explored the role of circulating miRNAs in SLE patients and non-pregnant lupus nephritis [8, 9], and some research has focused on circulating or placental EVs/miRNAs in pregnancy complications or SLE pregnancy [10, 11], the profile of urinary exosomal miRNAs specifically in pregnant women with SLE, especially when compared to healthy pregnant women, remains less comprehensively studied. Understanding the specific changes in the urinary exosomal miRNA cargo during late gestation in pSLE could reveal novel insights into the underlying pathological mechanisms and identify potential non-invasive biomarkers unique to this high-risk population.

Therefore, the aim of this study was to investigate the profile of urinary exosomal miRNAs in pregnant women with SLE during late gestation compared to healthy pregnant controls. We utilized high-throughput miRNA sequencing to identify differentially expressed miRNAs in urinary exosomes, followed by bioinformatics analysis, including target gene prediction and functional enrichment analysis, to elucidate the potential biological roles and pathways associated with the observed miRNA alterations in pSLE. This study seeks to contribute to the identification of potential non-invasive biomarkers and therapeutic targets for better management of pregnancy in women with SLE.

## 2. Materials and methods

### 2.1 Study Design and Participants

This study employed a comparative approach to analyze the profile of urinary miRNAs in pregnant women with SLE compared to healthy pregnant controls. A total of 11 pregnant women were enrolled, including 6 patients diagnosed with SLE and 5 healthy pregnant women. All participants were recruited from the Department of Obstetrics at Qilu Hospital of Shandong University between November 2022 and October 2023.

SLE patients were diagnosed based on the 2019 European League Against Rheumatism/American College of Rheumatology Classification Criteria for SLE [12]. Healthy pregnant women had no known history of SLE or other autoimmune diseases and were experiencing uncomplicated pregnancies and deliveries.

Ethical approval for this study was obtained from the Ethics Committee of Qilu Hospital of Shandong University (Ethics Number: KYLL-202210-046), and written informed consent was obtained from all participants prior to their enrollment. Participants with other concurrent severe medical conditions, active infections in the pregnant period, or other complications unrelated to SLE that might affect urinary exosome composition were excluded.

### 2.2 Clinical Data Collection

Relevant clinical data were collected for all participants during the late gestation period. For SLE patients, this included demographic information (age), obstetric history (parity, gestational age at recruitment and delivery), history of SLE diagnosis, SLE disease activity during pregnancy, current medications, presence of lupus nephritis during pregnancy, proteinuria levels, and other relevant antenatal clinical parameters. For healthy controls, demographic information, obstetric history, and confirmation of no significant medical history or antenatal complications were recorded. Pregnancy outcomes were also collected retrospectively for both groups. For research purposes, we accessed these clinical data on May 19, 2025, and authors had access to information that could identify individual participants during or after data collection.

### 2.3 Urine Sample Collection and Exosome Isolation

A clean catch mid-stream urine sample 50 mL was collected from each participant prior to delivery. Samples were immediately stored at 4°C and processed within 4 hours of collection to prevent degradation. For exosome isolation, urine samples were first centrifuged at 2,000×g for 10 minutes at 4°C to remove cellular debris. The supernatant was then processed for exosome isolation using sequential ultracentrifugation: 10,000 ×g for 30 minutes to remove microvesicles, followed by 100,000×g for 70 minutes at 4 ° C. The pellet was washed with sterile PBS and a final pellet was obtained by re-centrifugation at 100,000×g for 70 minutes. The exosome pellet was then resuspended in a suitable volume of PBS or lysis buffer. The identification of isolated exosomes was conducted by transmission electron microscopy (TEM, JEM-1200EX, Japan) and Nanoparticle Tracking Analyzer (NTA, Zetaview, Germany), and the protein concentration was quantified using the Micro BCA protein analysis kit (Boster, China). The expression of characteristic proteins TSG101 (1:1000; ab125011, Abcam) and CD81 (1:1000; 52892, CST) were detected by Western blot (WB).

### 2.4 Exosomal miRNA Extraction and miRNA Sequencing

Total miRNA was extracted from the isolated urinary exosomes using the miRcute miRNA Isolation Kit (TIANGEN, China) following the manufacturer’s protocol. Exosomal miRNA samples were prepared for small RNA sequencing. Differentially expressed miRNA were screened using DESeq2, with │ log2fold change │ ≥ 1 and adjusted *P* <0.05 as criteria. All RNA sequencing samples were commissioned by Xiuyue Biol (China).

### 2.5 Target Gene Prediction

Putative target genes for the significantly differentially expressed miRNAs identified in the previous step were predicted using several established online prediction databases/tools, including TargetScan [13], miRDB [14], miEAA[15], miRTarBase [16].

### 2.6 Functional Enrichment Analysis

Functional enrichment analyses, including Gene Ontology (GO) and KEGG pathway analysis, were performed on the compiled list of predicted and/or validated target genes of the differentially expressed miRNAs using clusterProfiler R package. Terms or pathways with an adjusted *P*-value < 0.05 were considered significantly enriched.

### 2.7 Statistical Analysis

Statistical analysis was conducted using GraphPad Prism 8.0.2 (GraphPad Software, USA). Quantitative data were first evaluated for normal distribution and reported as the mean ± standard error, followed by a variance homogeneity test. The t-test or one-way ANOVA was employed to ascertain statistically significant differences between two or among multiple homogeneous groups. For significant ANOVA outcomes, Tukey’s multiple comparison test and Dunnett’s multiple comparison tests were applied to analyze the differences between these groups. Results with *P* values less than 0.05 were considered statistically significant.

## 3 Results

### 3.1 Clinical Characteristics of Participants

In this study, a total of 11 pregnant women were enrolled and divided into two groups: 5 healthy pregnant controls and 6 pregnant women with SLE (pSLE). The demographic and clinical characteristics of these participants are summarized in Table 1. Comparison of clinical variables between the two groups revealed significant differences in levels of alanine aminotransferase (ALT) and aspartate aminotransferase (AST). ALT levels were significantly higher in the pSLE group compared to controls (22.33±11.52 U/L vs. 8.48±3.11 U/L, p = 0.0310), as were AST levels (36±15.75 U/L vs. 17.96±5.49 U/L, p = 0.0373).

**Table 1.** Clinical Characteristics of Participants.

| Variables | Control | pSLE | *P*-value |
| --- | --- | --- | --- |
| Age (years) | 32.6±2.51 | 29.33±3.14 | 0.0875 |
| WBC (×10<sup>9</sup>/L) | 7.878±1.88 | 7.49±1.56 | 0.7223 |
| Hb (g/L) | 112.2±13.37 | 115.5±20.13 | 0.7531 |
| PLT (×10<sup>9</sup>/L) | 233.8±55.51 | 205.8±32.81 | 0.3583 |
| ALT (U/L) | 8.48±3.11 | 22.33±11.52 | 0.0310* |
| AST (U/L) | 17.96±5.49 | 36±15.75 | 0.0373* |
| TBIL (μmol/L) | 8.12±6.42 | 9.02±5.01 | 0.8112 |
| TP (g/L) | 62.6±2.93 | 68.35±6.08 | 0.0773 |
| ALB (g/L) | 34.6±2.60 | 35.97±3.85 | 0.5027 |
| Crea (μmol/L) | 36±9.43 | 36(33.5,69) | 0.3052 |
| Urea (mmol/L) | 2.64±0.59 | 3.75 (2.8,6.23) | 0.1255 |
| Proteinuria positive | 0/5 (0%) | 2/6 (33.3%) | 0.4545 |
| Preterm | 0/5 (0%) | 3/6 (50%) | 0.1818 |

No statistically significant differences were observed for other measured continuous variables, including age (p = 0.0875), white blood cell (WBC) count (p = 0.7223), hemoglobin (Hb, p = 0.7531), platelet (PLT) count (p = 0.3583), total bilirubin (TBIL, p = 0.8112), total protein (TP, p = 0.0773), albumin (ALB, p = 0.5027), creatinine (Crea, p = 0.3052), and urea levels (p = 0.1255).

### 3.2 Isolation and Identification of Urine Exosomes

The overall experimental workflow is illustrated in Figure 1A. TEM revealed vesicles with typical morphology (Fig. 1B). WB confirmed the presence of canonical protein markers TSG101, CD81 (Fig. 1C). NTA showed a population of particles within the expected exosome size range and concentration (Fig. 1D). Collectively, these findings validated the successful isolation and characterization of exosomes from urine samples.

**Figure 1.**
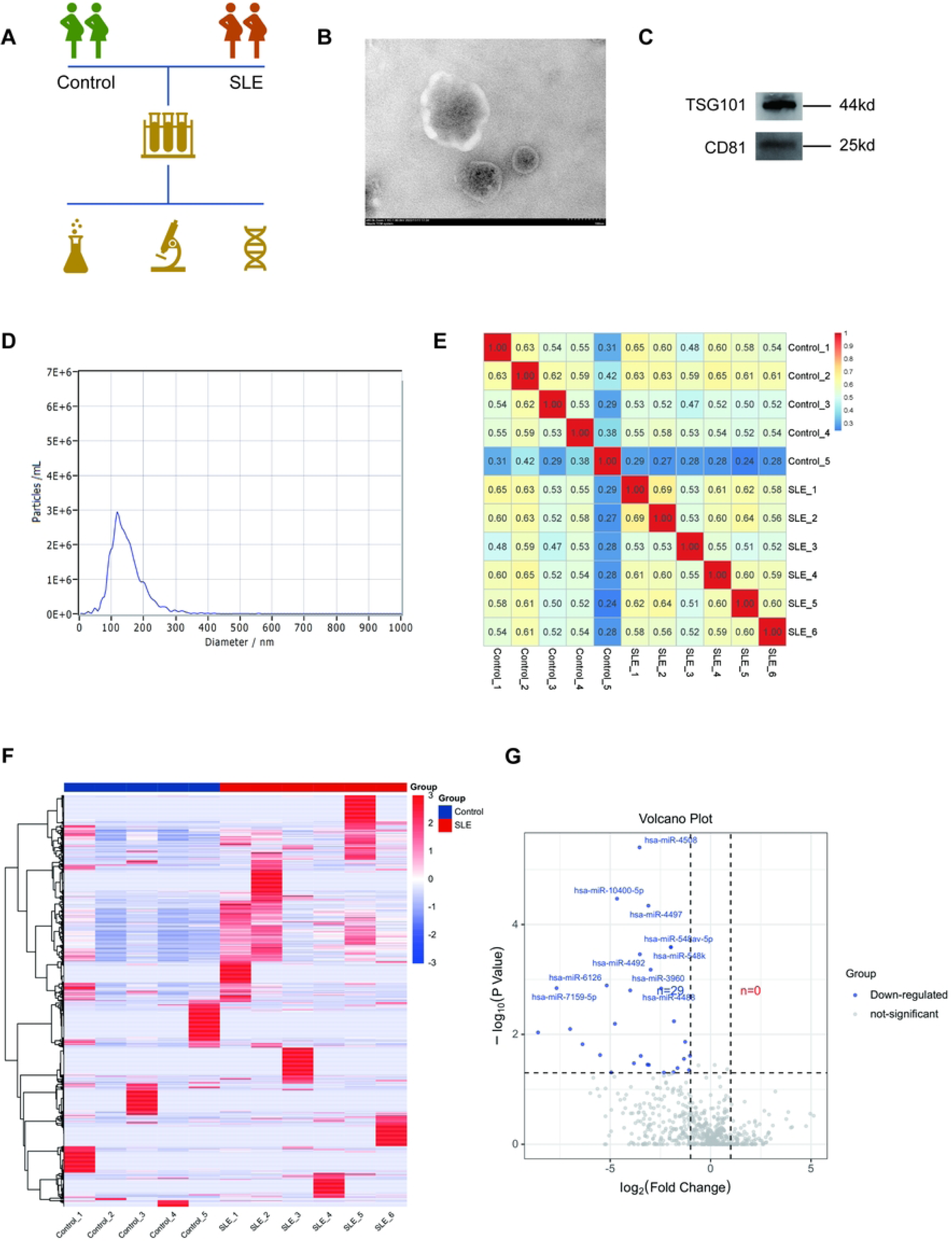
Isolation, Characterization, and Differential miRNA Analysis of Urine Exosomes. A. Schematic illustration of the overall experimental workflow. B. Representative TEM image showing the characteristic morphology of isolated urine exosomes, displaying their typical cup-shaped or spherical bilayer membrane structure. C. WB analysis confirming the presence of canonical exosomal protein markers, including TSG101 and CD81, in the isolated urine exosomes. D. NTA graph illustrating the size distribution and concentration of isolated particles, consistent with the expected size range of exosomes. E. Pearson correlation heatmap demonstrating the global relationships among all sequenced miRNA samples. F. Heatmap displaying the expression profiles of the significantly differentially expressed miRNAs identified between the control and disease groups across all samples. G. Volcano plot visualizing the results of the differential miRNA expression analysis.

### 3.3 Urinary Exosome miRNA Sequencing and Differential Analysis

Figure 1E shows a Pearson correlation heatmap of all sequencing samples, indicating good separation between the control and disease groups and high homogeneity within each group. Figure 1F displays a heatmap visualizing the expression patterns of the differentially expressed miRNAs. Further analysis using a volcano plot (Fig. 1G) identified a total of 29 significantly downregulated miRNAs. Among these, has-miR-4508, has-miR-10400-5p, and has-miR-4497 were the top three most significant based on their *P*-values.

### 3.4 Enrichment Analysis of Differentially Expressed miRNAs and Target Genes

Direct KEGG enrichment analysis of the differentially expressed miRNAs was performed using the miEAA tool. This analysis revealed significant enrichment in various metabolic pathways, including Cysteine and methionine metabolism, Glycolysis/Gluconeogenesis, Biosynthesis of unsaturated fatty acids, and Fatty acid elongation. Furthermore, several immune-related pathways, such as Th1 and Th2 cell differentiation, the Toll-like receptor signaling pathway, and the NOD-like receptor signaling pathway, were also identified (Fig. 2A).

**Figure 2.**
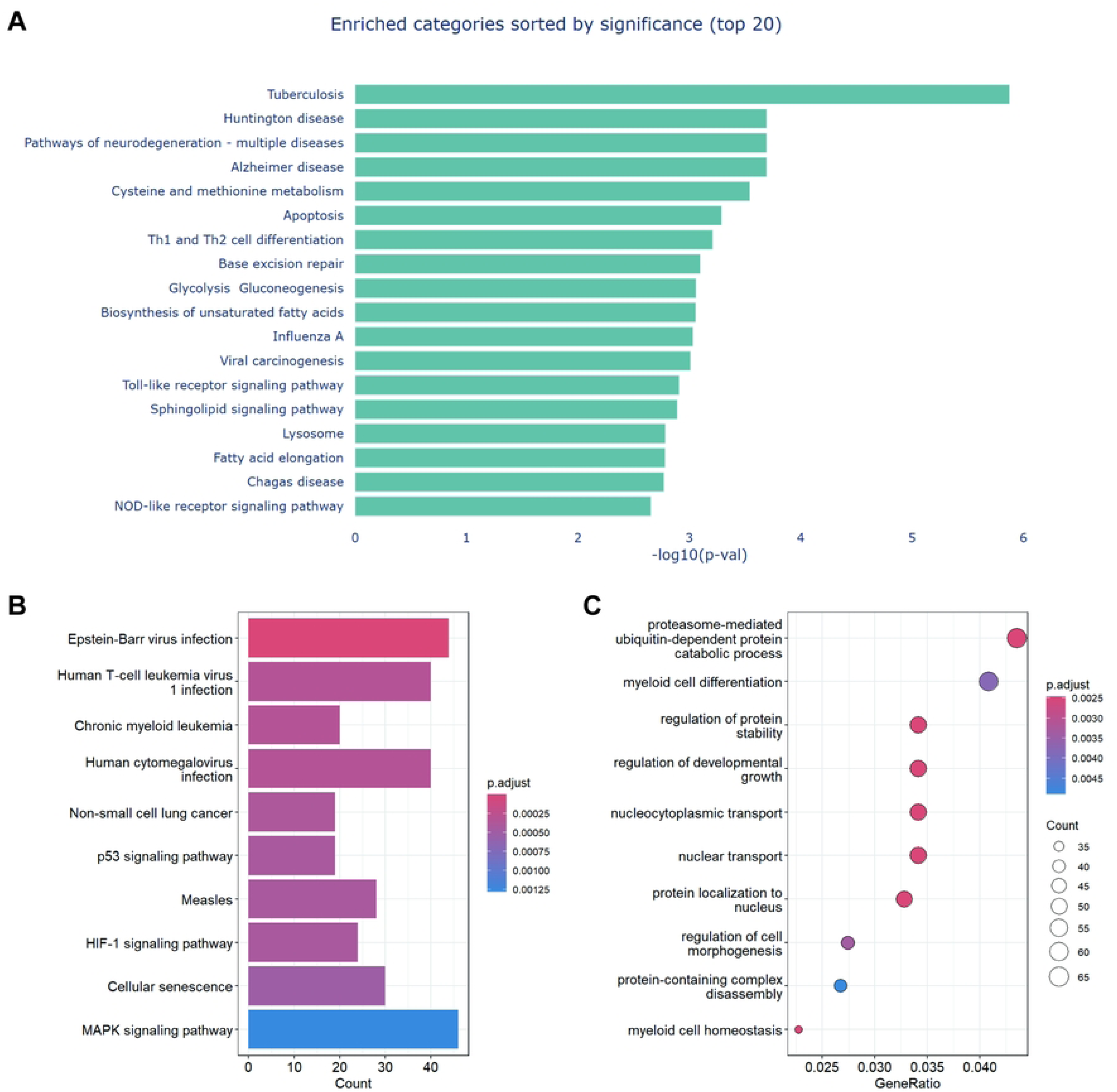
Functional Enrichment Analysis of Differentially Expressed miRNAs and Their Experimentally Validated Target Genes. A. KEGG pathway enrichment analysis of the significantly differentially expressed miRNAs, performed using the miEAA tool. B. KEGG pathway enrichment analysis of the experimentally validated target genes of the differentially expressed miRNAs, retrieved from miRTarBase. C. GO enrichment analysis of the experimentally validated target genes.

To investigate the functional implications of these miRNAs, experimentally validated target genes were retrieved from miRTarBase. Subsequent KEGG enrichment analysis of this comprehensive set of target genes highlighted pathways relevant to pregnancy outcomes in the pSLE group, specifically showing enrichment in virus infection pathways such as Epstein-Barr virus infection and Human cytomegalovirus infection. Additionally, pathways associated with cellular senescence, including the p53 signaling pathway and the broad category of Cellular senescence, were significantly enriched (Fig. 2B). Complementary Gene Ontology (GO) enrichment analysis of the same target gene set indicated involvement in biological processes related to protein stability and protein transport (Fig. 2C).

### 3.5 High-Level Pathway Enrichment Networks of Target Genes

To provide a higher-level overview of the biological functions represented by the enriched target gene pathways, an advanced analysis of these enrichment results was conducted. The KEGG enrichment findings indicated significant enrichment in various metabolic pathways, comprehensively covering carbohydrate, lipid, and amino acid metabolism (indicated by blue ellipses), alongside multiple immune-related pathways (indicated by orange ellipses) (Fig. 3A). Complementary Gene Ontology (GO) analysis of the pathways (or the target genes within these pathways) revealed that the target genes of the differentially expressed miRNAs are involved in biological processes related to mRNA and protein stability (pink ellipse), metabolic pathways under stress response (blue ellipse), and cell adhesion functions (purple ellipse) (Fig. 3B).

**Figure 3.**
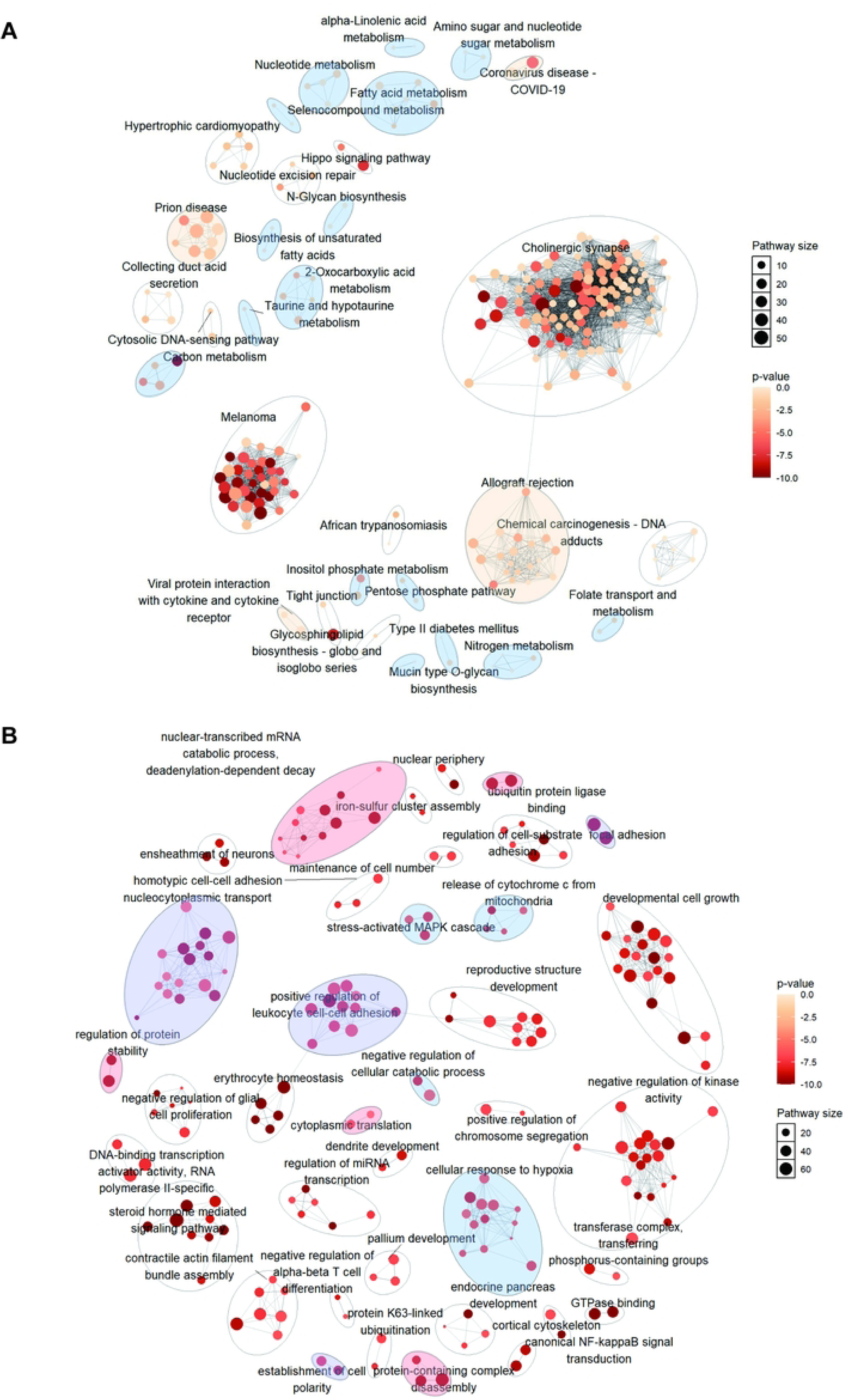
High-Level Functional Pathway Enrichment Networks of Differentially Expressed miRNA Target Genes. A. High-level KEGG pathway network analysis of the target genes, visualizing the broader functional categories of significantly enriched KEGG pathways. Blue ellipses represent metabolic pathway categories (carbohydrate, lipid, amino acid metabolism), and orange ellipses represent immune-related pathways. Connections indicate relationships or shared genes between pathways. B. High-level GO network analysis of the target genes or their associated enriched pathways, depicting major biological process categories. The pink ellipse represents processes related to mRNA and protein stability, the blue ellipse represents metabolic pathways under stress response, and the purple ellipse represents cell adhesion functions. Connections indicate relationships or shared genes/processes.

### 3.6 Enrichment Analysis of Target Genes for the Top Three Differentially Expressed miRNAs

Enrichment analysis was performed on the predicted target genes of the top three miRNAs with the largest fold changes (hsa-miR-8063, hsa-miR-7159-5p, and hsa-miR-3178). Gene Ontology (GO) enrichment analysis indicated involvement in biological processes related to mRNA synthesis, processing, and stability (Fig. 4A). KEGG pathway enrichment analysis revealed significant enrichment in the AMPK signaling pathway and senescence-associated pathways (Fig. 4B).

**Figure 4.**
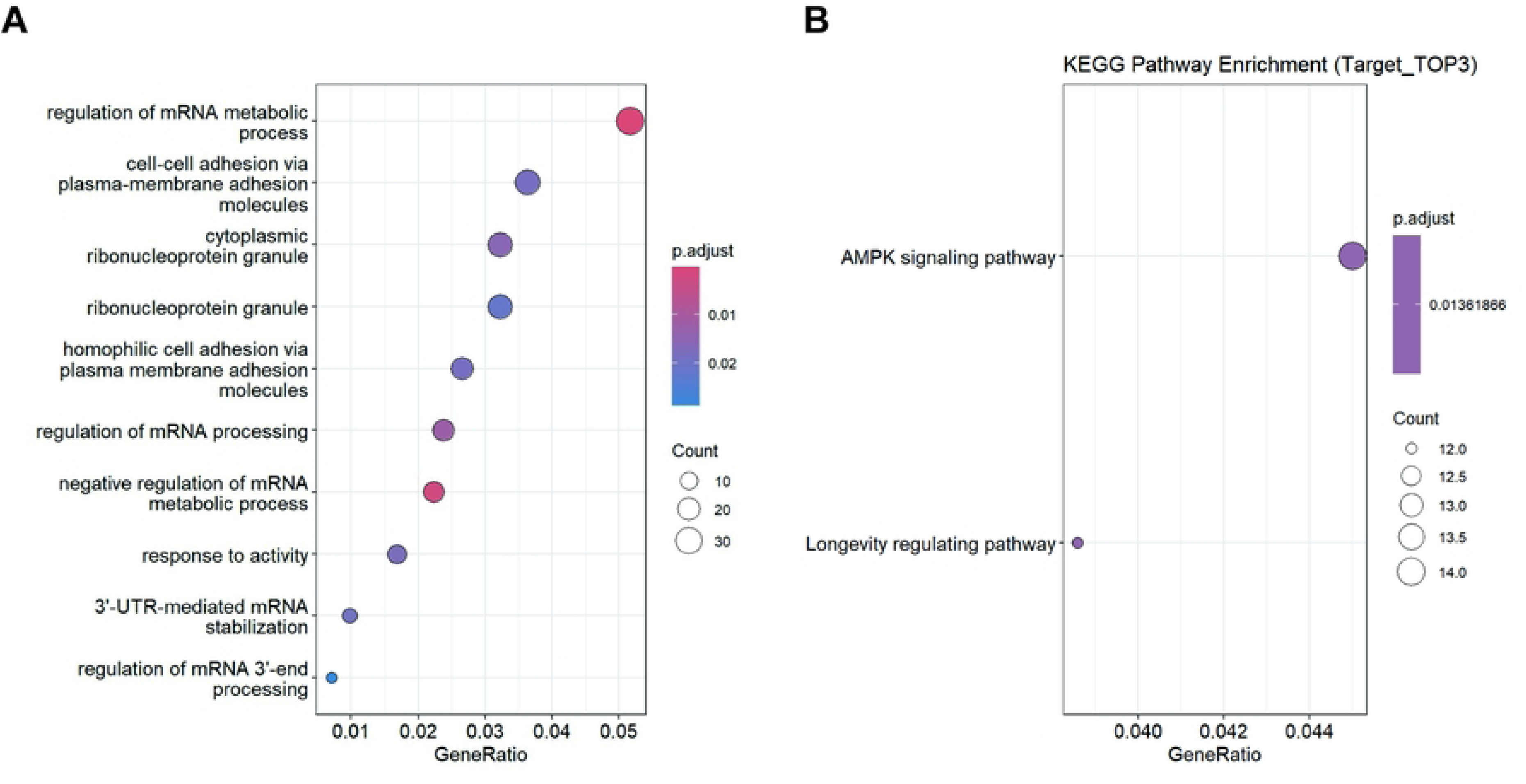
Functional Enrichment Analysis of Target Genes for the Top Three Differentially Expressed miRNAs. A. GO enrichment analysis of the predicted target genes for the three miRNAs with the largest fold changes. B. KEGG pathway enrichment analysis of the predicted target genes for the same top three miRNAs.

## 4 Discussion

pSLE presents significant challenges due to increased risks of both maternal and fetal complications. Non-invasive biomarkers capable of reflecting disease activity and predicting adverse outcomes are urgently needed for improved management of this high-risk population. EVs, particularly urinary exosomes, offer a promising source of such biomarkers, providing a window into systemic and renal pathological processes[17]. This study aimed to investigate the profile of urinary exosomal miRNAs in pregnant women with SLE during late gestation compared to healthy pregnant controls, and to explore their potential functional roles.

Our primary finding revealed a distinct profile of urinary exosomal miRNAs in late gestation pSLE, with 29 miRNAs being significantly downregulated compared to healthy pregnant controls. This differential expression suggests that the cargo of urinary exosomes is altered in pSLE, potentially reflecting changes in the biogenesis, content sorting, or cellular origin of these vesicles under disease conditions during pregnancy. While previous studies have explored circulating miRNAs in SLE or placental EVs in pregnancy complications[8, 18, 19], the specific landscape of urinary exosomal miRNAs in pSLE, particularly during the critical late gestation period, has been less defined. The identification of downregulated miRNAs aligns with some reports of altered miRNA profiles in other biofluids or tissues in SLE, although discrepancies exist, highlighting the tissue/fluid-specific and context-dependent nature of miRNA dysregulation.

Further functional enrichment analysis provided insights into the potential biological implications of the observed miRNA alterations. Direct KEGG analysis of the differentially expressed miRNAs pointed towards their involvement in various metabolic and immune-related pathways, including those related to biometabolism, as well as pathways central to immune cell differentiation and innate immunity. These findings are highly relevant to SLE pathogenesis, which is characterized by systemic inflammation and increasing evidence of metabolic dysregulation contributing to immune dysfunction [20–22]. The enrichment of metabolic pathways suggests that altered urinary exosomal miRNAs might originate from cells with perturbed metabolism or could influence metabolic processes in target cells within the urinary tract or systemically.

Importantly, analyzing the predicted and experimentally validated target genes of these miRNAs revealed enrichment in pathways with direct relevance to the complications seen in pSLE. Notably, KEGG analysis highlighted enrichment in virus infection pathways and cellular senescence-related pathways. SLE has long been linked to viral triggers, and viral reactivation can potentially occur or exert pathogenic effects during pregnancy [23, 24]. The enrichment of senescence pathways is particularly intriguing, as cellular senescence is increasingly recognized as a contributor to chronic inflammation, tissue dysfunction, and various pregnancy complications, including preeclampsia and placental insufficiency[25]. This suggests that aberrant urinary exosomal miRNAs in pSLE may influence host responses to viral challenges or contribute to premature cellular aging, which could impact the kidney, placenta, or other organs relevant to pregnancy outcomes. Complementary GO analysis further indicated roles for the target genes in protein stability and transport, fundamental cellular processes whose disruption can impact cellular function and stress responses.

Delving deeper into the higher-level functional categories reinforced these findings, showing comprehensive involvement in broad metabolic and immune functions as identified by KEGG networks. GO analysis of the pathways further associated target genes with mRNA/protein stability, stress-response metabolism, and cell adhesion, aligning with the idea that these miRNAs target core cellular maintenance and stress adaptation mechanisms, potentially disrupted in the complex inflammatory and metabolic environment of pSLE during pregnancy.

Analysis focusing specifically on the target genes of the top three miRNAs with the largest fold changes further supported these themes. Their predicted target genes were enriched in mRNA synthesis, processing, and stability, underscoring the potential impact on fundamental gene expression regulation. KEGG analysis of these specific miRNAs’ targets highlighted the AMPK signaling pathway and senescence-associated pathways. The AMPK pathway is a central regulator of cellular energy homeostasis, metabolism, and stress response, and its dysregulation has been implicated in both SLE and senescence[26, 27]. This provides a compelling link between the observed metabolic, senescence, and core cellular function enrichments, suggesting that these top miRNAs might act through pathways connecting energy status, cellular aging, and inflammatory responses.

This study benefits from its non-invasive approach using urinary exosomes and its specific focus on the understudied pSLE population in late gestation. However, limitations include the relatively small sample size, which necessitates validation in larger, independent cohorts. The cross-sectional design provides a snapshot but does not allow for longitudinal monitoring of miRNA changes throughout pregnancy or in response to treatment. Furthermore, the bioinformatics prediction of target genes and pathways requires experimental validation to confirm direct interactions and functional consequences.

## 5 Conclusion

In conclusion, this study provides initial evidence that urinary exosomal miRNAs are significantly altered in pregnant women with SLE during late gestation. Functional enrichment analysis suggests that the target genes of these miRNAs are involved in critical biological processes highly relevant to pSLE pathophysiology and pregnancy complications, including metabolic regulation, immune response, viral handling, cellular senescence, and fundamental cellular maintenance. These findings highlight the potential of urinary exosomal miRNAs as non-invasive biomarkers for monitoring disease status or predicting risks in pSLE and warrant further investigation for their mechanistic roles and clinical utility in managing this high-risk pregnancy population.

## Acknowledgments

We would like to thank Dr. Zhang Shule from the Department of Pediatrics, Affiliated Provincial Hospital of Shandong First Medical University, for providing assistance with data analysis.

## Ethics approval and consent to participate

This study was approved by the Research Ethics Committee of Qilu Hospital of Shandong University (Reference number: KYLL-2024(ZM)-1325).

## Consent for publication

Not applicable

## Availability of data and materials

The gene expression data of miRNA-seq data are deposited to the GEO with the dataset identified GSE298542.

## Authors’ contributions

Dong Li: Validation, Investigation, Data curation, Writing original draft. Anna Li: Validation, Investigation, Data curation, Formal analysis. Linghong Liu: Investigation, Data curation. Yuan Liu: Methodology, Writing-review & editing, Supervision, Project administration, Funding acquisition.

